# In vivo semen characterization and seasonal variation in *Procavia capensis*

**DOI:** 10.1101/2025.08.22.671636

**Authors:** Tal Raz, Hunter Alan Warick, Stav Asulin-Schnaiderman, Nuphar Shidlovsky, Nathalie Weizmann, Lee Koren

**Affiliations:** Koret School of Veterinary Medicine, Robert H. Smith Faculty of Agriculture, Food and Environment, The Hebrew University of Jerusalem, Rehovot, Israel; Advanced Academic Programs, Krieger School of Arts and Sciences, Johns Hopkins University, Baltimore, MD, USA; Faculty of Life Sciences, Bar Ilan University, Ramat Gan, Israel

**Keywords:** Spermatozoa, Sperm ultrastructure, Sperm morphology, Sperm morphometry, Reproductive seasonality, Electroejaculation, Rock hyrax, Wildlife reproductive biology

## Abstract

Seasonal reproduction is common among wild mammals, but male fertility traits are often understudied. The rock hyrax (*Procavia capensis*) is a seasonal breeder with a narrow reproductive window, yet its semen characteristics and seasonal variation remain poorly understood. Our objectives were to develop and validate an electroejaculation protocol for semen collection, and to enable, for the first time, in vivo assessments of sperm morphology, ultrastructure, morphometry, and seasonal variation in semen quality in both captive and wild populations. A total of 70 semen collection attempts were conducted: 17 in captive males at ∼monthly intervals over one year, and 53 in wild males just before peak mating season and again 2–4 weeks later, across three consecutive years (2021–2023). Electroejaculation was well tolerated and effective, particularly around the mating season, eliciting consistent responses and yielding sperm-containing ejaculates in 88.7% of wild procedures. Sperm morphometry revealed a mean total length of ∼56µm, with ultrastructural features resembling other eutherian mammals. Seasonal analysis demonstrated significantly higher sperm concentration, motility, and normal morphology during peak mating season compared to later samples. Post-peak samples showed increased structural abnormalities, including midpiece and principal piece defects, and signs of disrupted spermatogenesis and epididymal maturation. Cytology and high-resolution scanning electron microscopy confirmed these findings, revealing compromised sperm integrity and elevated round cell counts outside the breeding peak. This study establishes the first in vivo semen collection protocol and comprehensive semen evaluation in the rock hyrax, revealing seasonal variation in male fertility and enabling repeatable, non-lethal reproductive monitoring.

## Introduction

Many wild mammals exhibit pronounced seasonal variation in reproductive activity, yet male fertility traits remain poorly characterized in most species. The rock hyrax (*Procavia capensis*) is a medium-sized terrestrial mammal native to Africa and the Middle East, known for its complex social structure and distinct seasonal reproductive behavior (Bordes, et al. 2022, Goll, et al. 2023). In Israel, the mating season is brief and peaks in the summer (Bar Ziv, et al. 2016, Demartsev, et al. 2023, Koren, et al. 2006, 2008). The gestation period of the rock hyrax is approximately 7–8 months, with females giving birth to litters averaging three (range: 1–6) precocial young (Glover and Sale 1968, Sale 1965). The species has a zonary, hemochorial placenta, and both monozygotic and dizygotic twinning have been reported (Soma, et al. 1976). Male rock hyraxes typically reach sexual maturity at 16–17 months, and, unlike some other mammals, no distinct ejaculate fractions (e.g., gel plugs or honey fractions) have been described in this species.

Seasonal influences on male reproductive parameters are well documented in many domestic species, with notable changes in semen quality and fertility outcomes (Setchell 1993). However, the fundamental semen characteristics and their seasonal variation in rock hyraxes are poorly understood, limiting insights into their reproductive physiology and efforts related to conservation or population control. Anatomical studies based on deceased or euthanized animals have shown that, unlike most mammals, rock hyrax testes are located intra-abdominally, caudal to the kidneys, attached to the dorsal body wall (Glover 1973, Glover and Sale 1968). The ascrotal testes and accessory sex glands (prostate, seminal vesicles, and bulbourethral glands) undergo significant seasonal changes, enlarging during the breeding season and regressing afterward (Millar 1972, Millar and Glover 1970). The penis is vascular, and erection is achieved through the engorgement of blood within the cavernous tissue (Glover and Sale 1968). Previous research on male rock hyrax reproduction has primarily focused on behavior and steroid hormone levels in hair (Barocas, et al. 2011, Goll, et al. 2023, Koren, et al. 2008, Weissman, et al. 2019), with limited post-mortem assessments of reproductive anatomy or spermatozoa from the epididymis (Glover and Sale 1968, Millar 1972, Millar and Glover 1970). While informative, such invasive methods are impractical for longitudinal or conservation-based studies.

Electroejaculation, a non-terminal technique involving mild transrectal electrical stimulation to induce ejaculation, is commonly used in humans and various domestic and wild species (Abril-Sanchez, et al. 2019, Cary, et al. 2004, Chung, et al. 1998, Tecirlioglu, et al. 2002). Despite its widespread application, no established in vivo semen collection protocol for the rock hyrax has been reported, and data on sperm structure or seasonal trends are lacking.

Accordingly, the objectives of this study were to develop and validate a non-terminal, in vivo electroejaculation protocol for semen collection in male rock hyraxes and to characterize spermatozoa and semen parameters in both captive and wild populations. We further aimed to compare electroejaculation success and semen quality at different time points within the breeding season. To achieve this, we first conducted periodic evaluations in captive males across approximately one year; thereafter, we applied the protocol in desert field conditions, sampling wild males just before the peak mating season and again 3–4 weeks later, over three consecutive years. We hypothesized that electroejaculation could be effectively adapted for rock hyraxes, enabling repeatable semen collection and detailed analysis of sperm characteristics, with consistent seasonal patterns in semen quality detectable in both captive and wild males.

## Materials and methods

The study was approved by the Bar-Ilan University IACUC (023-b12061-25). Capture of wild hyraxes was authorized under permits issued and annually reviewed by the Israel Nature and Parks Authority. Captive males were handled and sampled from September 2017 to October 2018. For wild males, capture and semen collection attempts were conducted twice each year, during the summers of 2021–2023: once just before the peak breeding season (P-BS) and again approximately 3–4 weeks after (A-BS).

### Handling of captive male rock hyraxes

A captive rock hyrax colony was established at Bar-Ilan University (32°03’59.0“N, 34°50’34.4”E) in February 2017, with adult males (n=3; cM1-cM3, estimated to be 5–8 years old) and females (n=6) captured near Ein-Ya’akov in northern Israel (33°0′37″N, 35°13′45″E). Each individual was identified with a 5 g neck collar (Koren, et al. 2008). The colony was housed in a secure, fenced enclosure with ambient lighting and environmental enrichment (rocks, trees, and hiding spots) and was monitored primarily via video cameras to minimize disruption and stress. Animals were fed daily with fruits and vegetables, which were also routinely placed inside dedicated hyrax box traps to habituate them to trap entry (Koren, et al. 2006, 2008, Weissman, et al. 2019). Although maintained in captivity, the hyraxes were not habituated and could not be safely handled; therefore, traps were used as the safest and least stressful method for capture and sedation. Every 4–5 weeks, traps were activated before dawn for semen collection. Captured males were transferred (in covered traps) to a nearby lab room (∼5-minute walk) for semen collection, as detailed below. Electroejaculation procedures were limited to 10 minutes of rectal stimulation. Afterward, males were held in the traps for recovery (2–3 hours) before being returned to the colony.

### Handling of wild male rock hyraxes

Wild male hyraxes were captured at Ein Gedi Nature Reserve (31°28’17.0“N, 35°23’12.8”E) during the summers of 2021–2023. Capture and semen collection attempts occurred twice each year, just before the peak mating season (P-BS) and again 3–4 weeks later (A-BS). Each sampling session (P-BS; A-BS) lasted approximately 5–7 days. The mating season was predicted from years of colony monitoring and confirmed by daily observations during sampling years (Goll, et al. 2023, Koren, et al. 2006, 2008, Koresh, et al. 2016, Weissman, et al. 2019). Live box traps baited with cabbage and fruit were placed in natural crevices ∼90 minutes before dawn and checked every 1–2 hours until noon. Only mature males were anesthetized for sampling; females and juveniles were released. Most males were already marked with neck collars; unmarked individuals were collared during handling. To avoid repeated sampling, each male contributed only one semen sample per session. Electroejaculation was limited to < 20 minutes. Afterward, males recovered in shaded traps for 2–3 hours before being released at their capture site.

### Semen collection by electroejaculation

Captive males were anesthetized with intramuscular Ketamine (10 mg/kg) and Midazolam (5 mg/kg), while wild males received Ketamine only to ensure faster recovery in the field. Each animal was briefly examined to rule out illness or injury. Feces were cleared using diluted rectal lubricant (OB Lube, AgriLabs, MO, USA; 1:1 with water), administered 3–5 times in 5 mL volumes via the anus.

Semen collections were performed using a commercial electroejaculator system (Minitube GmbH, Germany) with a canine probe (22 cm total length; 6.35 cm stimulation length; 1.27 cm diameter). Animals were positioned laterally on their right side, and the lubricated probe was inserted rectally with the stimulation surface oriented ventrally.

Electrical stimulation was controlled manually in 8–10 second cycles, alternating in a ∼3:2 increase-to-decrease ratio. Voltage began at 0.5V and was raised in 0.5–1V increments every 2–6 cycles, up to a maximum of 10V, adjusted based on observed responses (e.g., muscle contractions, erection, and ejaculation), which were recorded. Once extruded, the penis was gently cleaned with wet 4×4 gauze and dried using KimWipes™ (Thermo Fisher Scientific, Israel). The penis was held to guide the glans into a 1.5 mL collection tube containing 500 µL of semen extender (EquiPlus, Minitube, Germany) pre-warmed to body temperature, and gently squeezed from base to tip to promote semen flow to the external urethral orifice. If a clear pre-sperm fraction was present (occasionally, in a tiny volume), it was wiped away. The raw ejaculate (typically a tiny drop of 1–5 µL) was collected by gently pressing the glans against the inner wall of the collection tube and mixing it immediately with the extender. Because the raw ejaculate volume was extremely small (1–5 µL) and could not be measured accurately without risking sample loss, the total sample volume used for all calculations was defined as 500 µL, corresponding to the fixed volume of extender; the ejaculate volume was therefore considered negligible relative to the extender. The resulting diluted semen preparation was defined as the semen sample and was used for all handling and analyses. Following the procedure, the probe and rectum were checked for signs of injury, and hyraxes were released back into their habitat only after they had fully recovered and were bright, alert, and responsive.

### Semen sample handling and evaluations

Immediately after collection, sperm presence and progressive motility were evaluated on wet mounts (two replicates per diluted semen sample) in accordance with the guidelines of the WHO Laboratory Manual for the Examination and Processing of Human Semen (World Health Organization 2021). A 10 µL well-mixed aliquot was placed onto a clean, pre-warmed microscope slide and carefully covered with a pre-warmed 22 mm × 22 mm coverslip. Motility was evaluated using a Nikon Eclipse E400 biological microscope equipped with phase-contrast optics. The examination was conducted by the same trained operator within each setting (S.A.S. for captive samples, H.W. for field samples), both of whom were trained and standardized by the first author (T.R.). The initial microscopic examination was performed using a ×10 objective (total magnification ×100) to obtain a quick overview of the diluted semen sample, ensure the absence of drift or fluid flow, and confirm that spermatozoa were evenly distributed across the slide. Motility was then assessed at ×400 total magnification (×40 objective) in approximately ten randomly selected fields, avoiding areas within 5 mm of the coverslip edge to prevent drying artefacts. The numbers of progressively motile spermatozoa (moving actively, either linearly or in large circles, covering a distance) and non-progressive or immotile spermatozoa were recorded and used to calculate the percentage of progressive motility.

Cytology smears were prepared using Nigrosin & Eosin (N&E; Hancock Stain, Animal Reproduction Systems, USA) as previously described (Brito, et al. 2010, 2011). Additional unstained smears were prepared, air-dried, and later stained with Diff-Quick (Medi-Market, Israel) for round cell identification, defined as cells lacking the typical characteristics of spermatozoa (head, midpiece, and tail), such as immature germ cells and leukocytes (Johanisson, et al. 2000). For sperm concentration analysis, 50 µL of extender-diluted semen sample was further diluted 1:10 in a saline-formalin solution (approximately 0.14 M formalin, equivalent to ∼0.4% formaldehyde) to immobilize and preserve the spermatozoa until counting (Brito, et al. 2016, World Health Organization 2021).

Microscopic evaluations of sperm morphology and concentration were performed in the lab after the completion of each study phase by a single experienced observer (T.R), who was blinded to sample identity (male, sampling session, date, etc.). On N&E-stained slides, at least 100 spermatozoa per sample were evaluated at ×1000 magnification using an oil immersion ×100 objective (Axio Imager M1 microscope, Zeiss, Germany). Assessed abnormalities included acrosome defects, head, midpiece, and principal piece anomalies, detached heads, and cytoplasmic droplets (Barth and Oko 1989). The round cell-to-sperm ratio was determined by counting all round cells and spermatozoa in successive microscopic fields until a total of at least 300 spermatozoa had been counted. Sperm concentration was measured using a Neubauer hemocytometer count under phase contrast microscopy with a ×40 objective on an Axio Imager M1 microscope (Brito, et al. 2016, World Health Organization 2021). Based on these data, the following parameters were calculated for each diluted semen sample:

𝑻𝒐𝒕𝒂𝒍 𝑺𝒑𝒆𝒓𝒎^𝟏^ = (𝑽𝒐𝒍𝒖𝒎𝒆) × (𝑪𝒐𝒏𝒄𝒆𝒏𝒕𝒓𝒂𝒕𝒊𝒐𝒏)
𝑻𝒐𝒕𝒂𝒍 𝑴𝒐𝒕𝒊𝒍𝒆 𝑺𝒑𝒆𝒓𝒎 = (𝑻𝒐𝒕𝒂𝒍 𝑺𝒑𝒆𝒓𝒎) × (%𝑴𝒐𝒕𝒊𝒍𝒊𝒕𝒚)
𝑻𝒐𝒕𝒂𝒍 𝑷𝑴𝑴𝑵^𝟐^ = (𝑻𝒐𝒕𝒂𝒍 𝑴𝒐𝒕𝒊𝒍𝒆 𝑺𝒑𝒆𝒓𝒎) × (%𝑵𝒐𝒓𝒎𝒂𝒍 𝑴𝒐𝒓𝒑𝒉𝒐𝒍𝒐𝒈𝒚)
^1^Calculated assuming a total diluted volume of 500 µL per semen sample.
^2^PMMN, Progressively Motile Morphologically Normal sperm cells per ejaculate.

For morphometry, N&E-stained sperm were imaged at 1000x (AxioCam HRc, Zeiss). ImageJ (NIH) was used to measure total sperm length, head length and width, and midpiece and principal piece lengths. Measurements were performed on 586 normal sperm from 56 images of three ejaculates (three males). Images were captured by one operator (T.R.), and morphometric measurements were performed by another (N.S.), who was blinded to image origin and sample identity.

### High-resolution scanning electron microscopy (SEM) of hyrax spermatozoa

Sperm samples were prepared for SEM by placing 20 µL of centrifuged, resuspended diluted semen sample onto poly-L-lysine-coated round glass coverslips (5mm diameter; Thermo Fisher Scientific Inc., Israel) in a 96-well plate, incubated for 1 hour in a humid chamber. Coverslips were pre-coated and air-dried. Samples were fixed with 4% PFA for 1 hour at room temperature, washed with PBS, and dehydrated through a graded ethanol series (20% to 100%). Dehydrated samples were dried using a critical point dryer (Quorum K850) with liquid CO₂, mounted on stubs, and gold-coated (Quorum Spatter Coater; Quorum Technologies, UK). Images were captured using a JEOL JSM-7800 high-resolution Scanning Electron microscope (JEOL Ltd., Japan) at 5 kV using secondary electron detection in high-vacuum mode.

### Statistical analyses

Statistical analyses were performed using Statistix 10 software (Analytical Software, FL, USA), and plots were generated with Prism 5.01 (GraphPad Software, CA, USA). Descriptive statistics were calculated for all semen parameters. All statistical tests were based on two-tailed hypotheses. The non-parametric Wilcoxon Rank Sum Test was used to compare two groups (e.g., P-BS vs. A-BS within a given year or summer vs. non-summer). Fisher’s Exact Test was applied to compare proportions (e.g., ejaculation success rates). Associations between continuous variables (e.g., sperm counts, motility) were assessed using Spearman’s Rank Correlation Coefficients (ρ). To evaluate seasonal variation while accounting for repeated sampling of the same individuals, Repeated Measures ANOVA was performed, followed by Tukey’s HSD All-Pairwise Comparisons test. This model included only males sampled in both P-BS and A-BS within the same breeding season. Male identity was included as a random factor (subject), with sampling session (P-BS/A-BS; within-subject factor), year (2021–2023; between-subject factor), and their interaction treated as fixed effects. P-values < 0.05 were considered statistically significant, while 0.05 ≤ P < 0.10 was noted as a trend. Results are presented as mean±SEM, unless indicated otherwise.

## Results

### Electroejaculation-based semen collection and parameters in captive rock hyrax males

From September 2017 to November 2018, a total of 17 electroejaculation attempts were conducted with three captive hyraxes (cM1-cM3), as detailed in Supplementary Table 1. Male cM3 was transferred in October 2017 to another colony due to circumstances unrelated to the study, and therefore, it was sampled only once. Capture success varied due to inconsistent trap entry, and trapping attempts were limited to minimize disruption and stress within the colony. Semen parameters from attempts that yielded sperm are shown in Table 1. All hyraxes recovered well without any adverse effects.

**Table 1.**
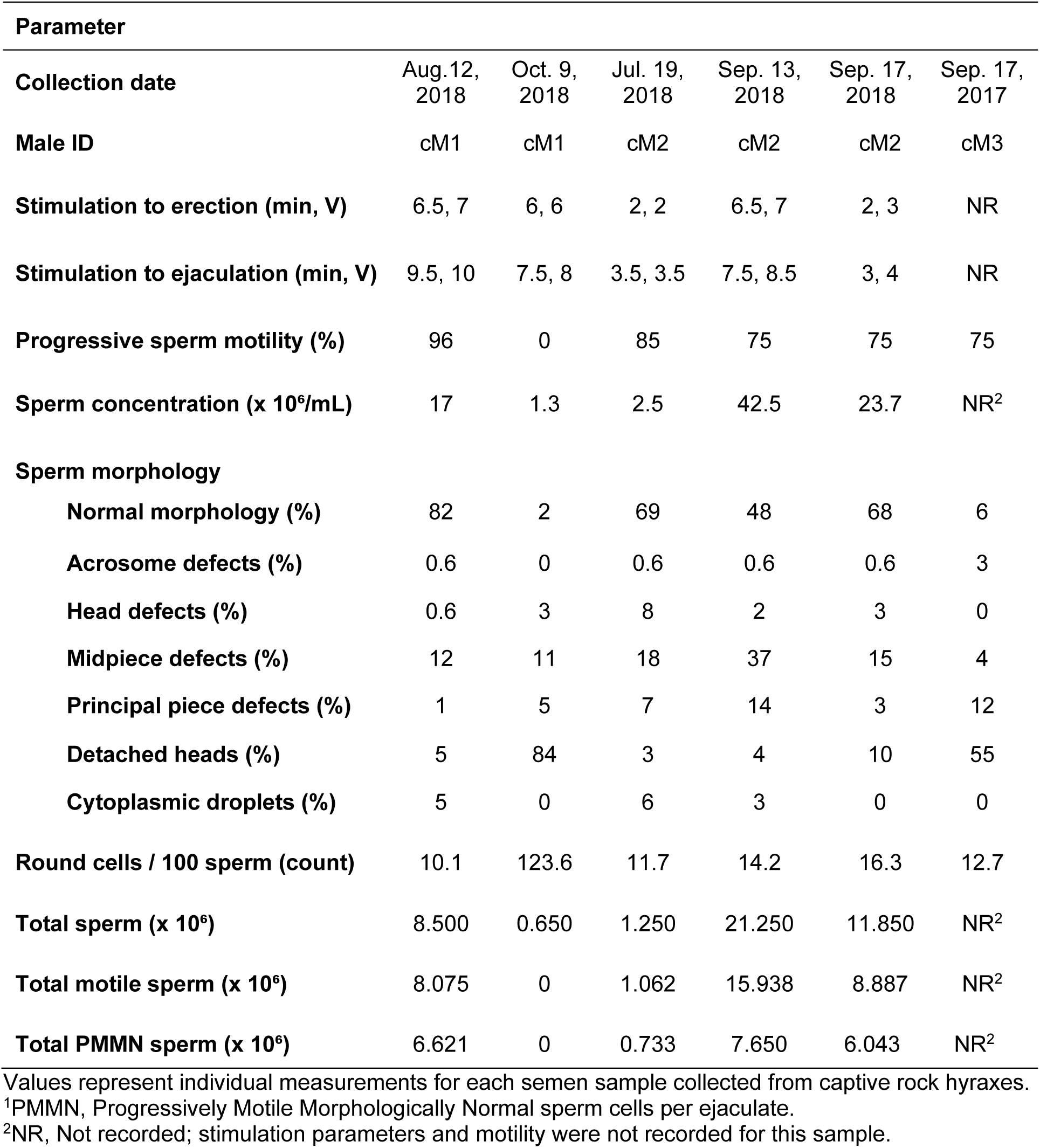
Semen characteristics of the six samples collected from captive rock hyraxes by electroejaculation.

Physical responses to electroejaculation stimulation included muscle contractions aligned with the electrical stimulation cycles, exhibiting strong extension of the limbs and neck as the stimulation increased, followed by relaxation as the stimulation decreased. Erection was achieved in all 17 attempts (100%), with an average induction time of 4.3±0.7 minutes (median: 3.25 minutes, range: 2–8.5 minutes; data recorded in 12 attempts) and peak stimulation of 4.8±0.65V (median: 4V, range: 3–8.5V; data recorded in 9 attempts). These parameters showed no significant difference between summer (defined from June 21st to September 21st) and non-summer months (Time, P=0.4889; Stimulation intensity, P=0.8095; Wilcoxon Rank Sum Test), or between attempts that resulted in sperm-containing samples and those that did not (Time, P=0.3699; Stimulation intensity, P=0.4524; Wilcoxon Rank Sum Test).

Ejaculation, defined as the release of fluid at the penile orifice, was observed in 70.6% of attempts (12/17), occurring exclusively from late spring through early autumn (April to early October), except one instance in January where only clear fluid without sperm cells was obtained (see Supplementary Table 1). The raw ejaculate volume was generally small (estimated as 1–5 µL), with slightly larger amounts recorded in two cases from August and September. The appearance of the fluid ranged from clear to milky opaque; only milky opaque samples contained sperm, while clear or clear-yellowish fluids lacked sperm (6/6, 100% vs. 0/6, 0%; P=0.0022). Two samples from May contained only round cells. There was no significant difference in the overall rate of fluid ejaculation between summer and the rest of the year (5/5, 100% vs. 7/12, 58.3%; P=0.2445), but milky opaque ejaculates containing sperm were significantly more likely to occur in summer (5/5, 100% vs. 1/12, 8.3%; P=0.0010).

As shown in Table 1, semen parameters from six sperm-containing diluted semen samples collected from captive rock hyraxes exhibited significant variation in motility, concentration, and morphology. Morphological assessment revealed a wide range in the percentage of normal sperm, while abnormalities in the midpiece, principal piece, and detached heads were common in lower-quality samples, typically collected after the breeding season. Notably, the two samples with the lowest percentages of normal morphology (collected in September 2017 and October 2018) showed a high proportion of detached heads (55% and 84%, respectively), consistent with sperm senescence typically observed during the non-breeding period (Koziol and Armstrong 2022, Okano, et al. 2009)

The number of round cells per 100 spermatozoa varied across diluted semen samples, ranging from 10.1 to 16.3 in most samples. A markedly elevated count was observed in one ejaculate from male cM1 (123.6 round cells/100 sperm), which was collected in October, well outside the typical breeding season. Spearman rank correlation analyses revealed that round cell abundance was negatively correlated with both progressive motility (ρ=–0.9411, P=0.0167) and normal morphology (ρ=–0.7714, P=0.1028), suggesting that an increased presence of round cells may be associated with reduced semen quality. Furthermore, the time elapsed since the peak breeding season (defined as mid-July) was positively correlated with round cell count (ρ=0.8407, P=0.0333) but negatively correlated with progressive motility (ρ=–0.8933, P=0.0333) and normal morphology (ρ=–0.8407, P=0.0583), supporting a temporal decline in semen quality in captive males as the season progressed.

### Sperm morphology and structural characteristics in rock hyraxes

Microscopic evaluation of N&E-stained hyrax semen cytology revealed a diverse spectrum of sperm morphologies, including both normal and abnormal forms (Fig. 1A–F), as well as round cells (Fig. 1G). Diff-Quick staining (Fig. 1H–J) enabled better identification of round cell types, including presumptive spermatocytes, spermatids, and degenerative forms such as medusa cells. High-resolution SEM imaging provided further insights into the ultrastructure of normal and abnormal hyrax spermatozoa, as illustrated in Fig. 2.

**Fig. 1.**
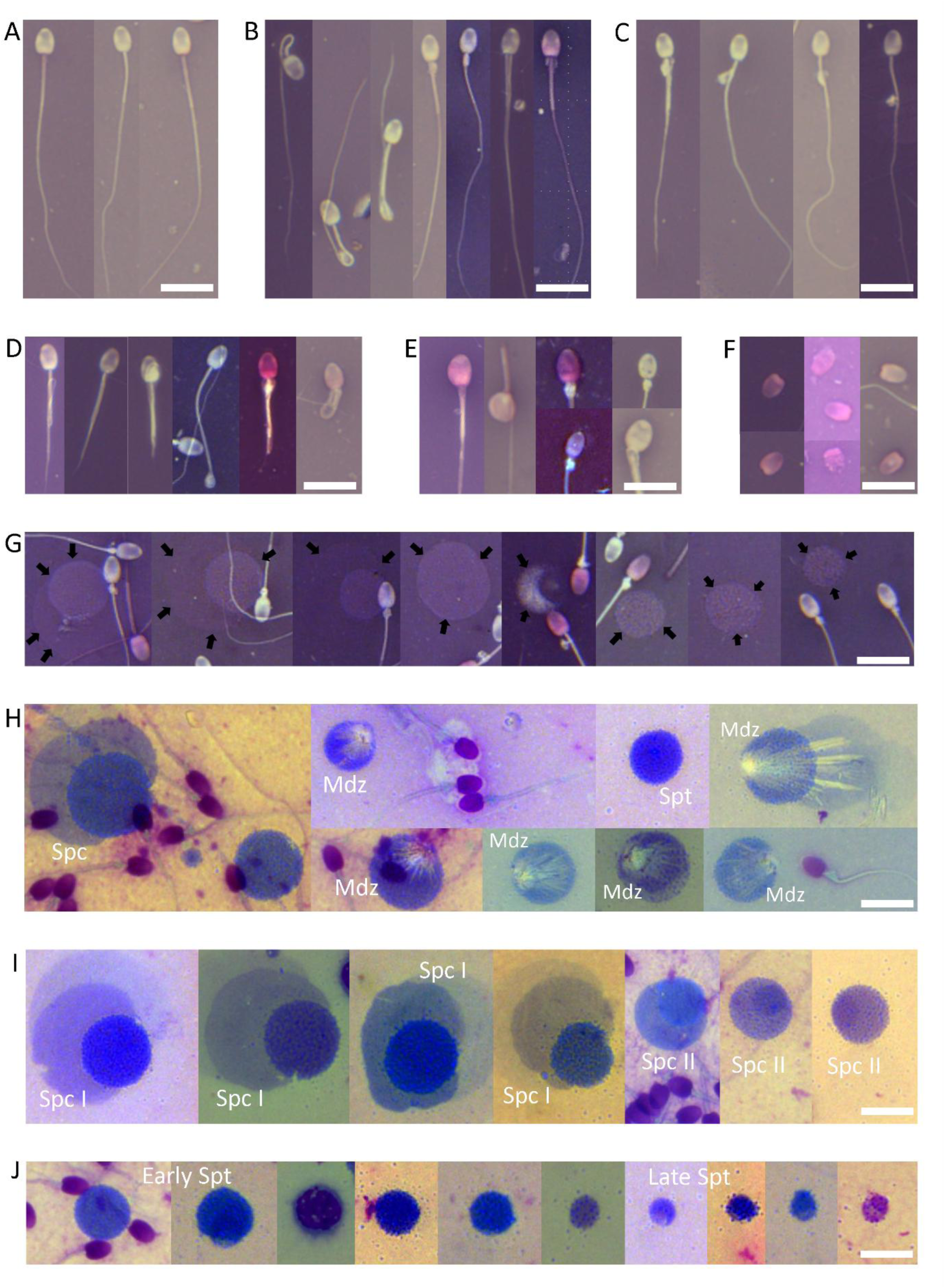
Microscopic imaging of rock hyrax semen cytology: spermatozoa and round cells. Panels A–G show Nigrosin & Eosin (N&E)–stained slides; Panels H–J show Diff-Quick–stained slides. All images are presented at the same magnification, with a scale bar representing 10 µm. Panels A–F present representative examples of individual spermatozoa and common morphological abnormalities, while Panels G–J focus on round cells. (A) Normal sperm. (B) Midpiece defects, including bowed midpiece, distal midpiece reflex, mitochondrial sheath defects, pseudodroplets, and accessory/multiple tails. (C) Cytoplasmic droplets, from proximal to distal locations. (D) Principal piece defects, including coiled tails, truncated tails, and Dag-like defects, characterized by strong coiling and folding of the tail (right image). (E) Head defects, including macrocephalic and microcephalic forms, vacuolated heads (craters), clumped chromatin, and ruffled acrosomes. (F) Detached sperm heads, many displaying additional head abnormalities as in panel E. (G) N&E-stained field with round cells (marked with arrows) intermixed with spermatozoa; note that round cell classification is limited with this stain. (H) Diff-Quick–stained field showing a mixture of round cells, such as presumed spermatocytes (Spc), round spermatids (Spt), and various medusa cells (Mdz); medusa cells are degenerating germ cells characterized by a central nucleus and radiating tail-like projections, typically indicative of abnormal spermatogenesis or post-testicular degradation. Note the improved cytological detail compared to N&E. (I) Presumptive primary spermatocytes (Spc I), large cells (∼12–18 µm) with pale cytoplasm and coarse chromatin, and secondary spermatocytes (Spc II), smaller (∼9–12 µm), with more condensed chromatin. (J) Presumed round spermatids in different developmental stages, from early (left) to late (right); early spermatids are small, round, and have uniformly dense chromatin, while late spermatids show slightly smaller size and chromatin condensation with initial polarization.

**Fig. 2.**
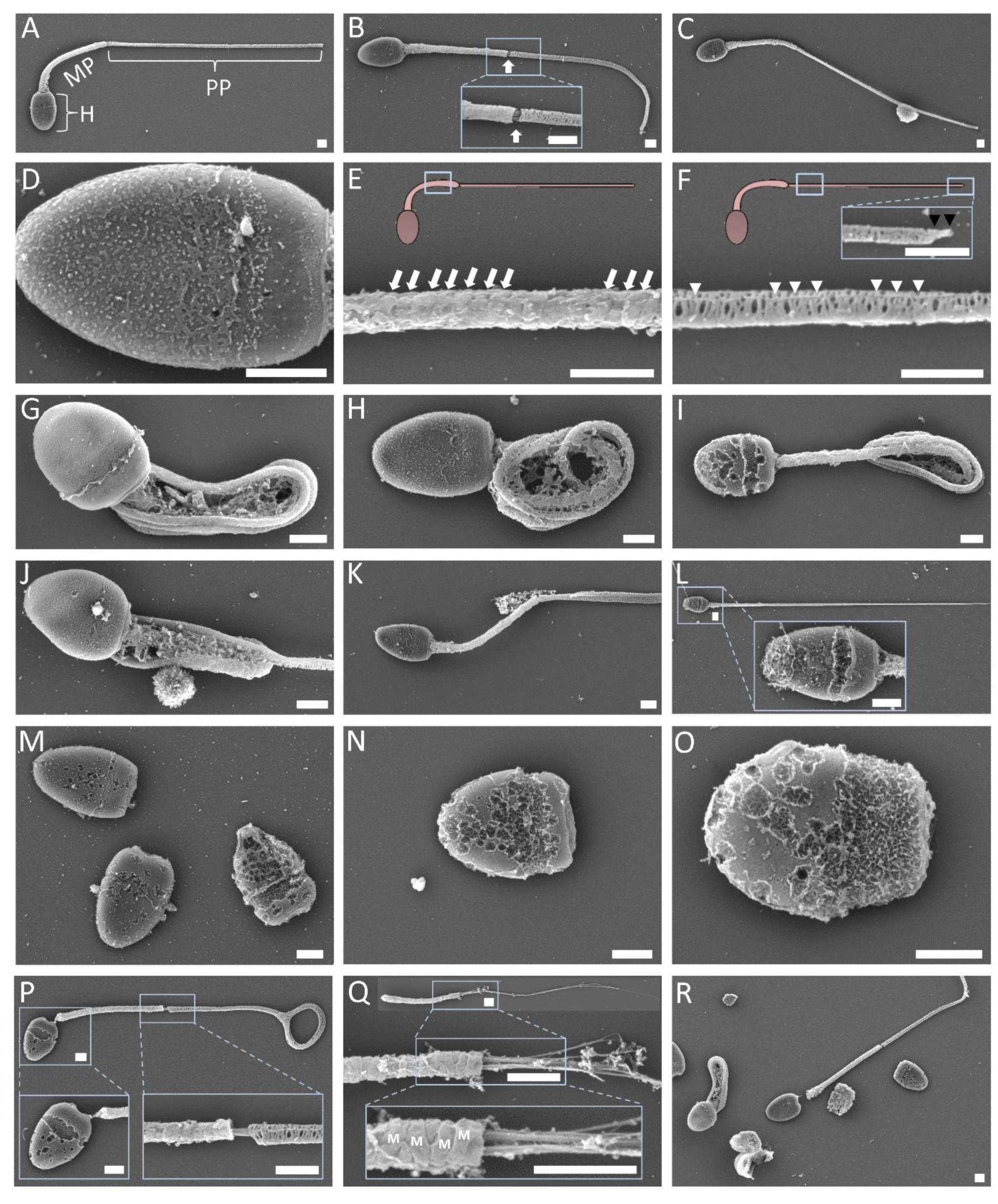
High-resolution scanning electron microscopy (SEM) of normal and abnormal rock hyrax spermatozoa. Micrographs were taken at various magnifications; scale bars represent 1 µm. (A) Intact spermatozoon with normal morphology, highlighting the head (H), midpiece (MP), and principal piece (PP). (B) Sperm with normal appearance but a torn fibrous sheath at the annulus (arrow), marking the junction between the MP and PP; note the slight exposure of axonemal microtubules in the high-magnification inset (lower box). (C) Sperm with a small cytoplasmic remnant located approximately two-thirds along the principal piece. (D) High magnification of a sperm head, showing intact plasma membrane over the acrosomal (apical) and post-acrosomal regions; the equatorial segment is evident. (E) An enlarged view of the midpiece (region boxed in sperm diagram) shows helically arranged mitochondria (indicated by white arrows). (F) High magnification of proximal PP (region boxed on the left in sperm diagram), displaying circumferential fibrous sheath ribs and longitudinal elements (white arrowheads). The small endpiece is visible at the top right inset and marked with black arrowheads. (G–H) Spermatozoa exhibiting strong coiling of the tail, consistent with Dag-like defects, often accompanied by a retained cytoplasmic droplet. (I) Spermatozoon showing a distal midpiece reflex anomaly, with a cytoplasmic droplet located within the coiled tail. In (G), the equatorial segment appears as a circumferential ridge, potentially due to mechanical stress, fixation artifacts, or early acrosomal reaction. In (I), the sperm head shows a porous, irregular surface, consistent with loss of the acrosomal cap and nuclear membrane compromise. (J) Sperm with a large proximal cytoplasmic droplet. (K) Sperm with a small distal cytoplasmic droplet and possibly a coiled PP, partially visible. (L) Longitudinal view of sperm with an apparently intact flagellum but missing the acrosomal cap and exhibiting membrane disruption (see zoomed-in head inset). (M–O) Detached heads showing progressive structural degradation, likely due to post-testicular aging, handling, or pathology, with acrosomal loss, membrane rupture, and chromatin exposure. (P) Severely degenerated spermatozoon with head degradation, partial detachment, disorganized midpiece (mitochondrial sheath disruption), and coiled PP. In the lower right inset, exposed axonemal microtubules are visible at the MP–PP junction (arrow). (Q) Detached, damaged sperm flagellum. The overview (top) and higher-magnification panels (middle and bottom) show the mitochondrial helical structure (M) and a few exposed axonemal microtubules, indicating the loss of membrane integrity. (R) Multiple spermatozoa and heads in various advanced degenerative stages, including distal midpiece reflex, complete head-tail separation, membrane loss, and structural disintegration.

Normal sperm morphometry results are presented in Table 2. No significant differences were found among males. The average total sperm length was 56.12±0.13 µm, with the principal piece comprising 72.57% and the midpiece 18.41% of the total length. When considering only the flagellum (midpiece plus principal piece), the midpiece accounted for 20.25% and the principal piece 79.75%. These values emphasize the elongated structure of rock hyrax spermatozoa and the distinct proportions of their components.

**Table 2.** Morphometric measurements and relative proportions of sperm structures in rock hyrax semen.

| Sperm parameter | Measurement ($\mu\text{m}$)<sup>1</sup> | Proportion of total sperm length (%)<sup>1</sup> |
| --- | --- | --- |
| Head Width | 3.49±0.01 (3.47–3.51) | 6.25±0.03 (6.02–6.03) |
| Head Length | 5.05±0.01 (5.03–5.07) | 9.03±0.03 (8.96–9.09) |
| Midpiece Length | 10.28±0.03 (10.22–10.33) | 18.41±0.09 (18.23–18.59) |
| Principal Piece Length | 40.80±0.13 (40.54–41.06) | 72.57±0.11 (72.34–72.79) |
| Total Sperm Length | 56.12±0.13 (55.88–56.38) | 100.00 |

<sup>1</sup> Values are presented as mean±SEM, with the 95% confidence interval shown in parentheses.

### Electroejaculation-based semen collection and parameters in wild rock hyrax males – Summer of 2021

During the summer of 2021, 17 electroejaculation procedures were performed in the field: 12 just before the peak mating season (P-BS; July 5–14, n=9 males) and 5 during the late season (A-BS; August 2–8, n=5 males). Erection was achieved in all cases (100%), and sperm cells were collected and confirmed in 14 of 17 ejaculates (82.4%) within a mean stimulation of 11.2±1.1 minutes (Median 10 minutes; range 5–20 min). In the P-BS session, sperm cells were identified in 9 of 12 electroejaculation attempts (75%), each from a different male. One male required three attempts, with only clear pre-sperm fluid collected in the first two before sperm was obtained on the third attempt. Another male yielded only pre-sperm fluid initially but produced a semen sample a few days later in the same session. In A-BS, all five procedures yielded sperm-containing samples (100%).

Analyses of semen parameters from all 14 diluted semen samples collected during the summer of 2021 revealed considerable variability in sperm quantity and quality. There were no significant differences in body weight (P-BS: 2.7±0.2 kg vs. A-BS: 2.5±0.2 kg; P=0.5000) or collection duration (12.1±1.5 min vs. 10.0±1.1 min; P=0.2792) between P-BS and A-BS samples. However, as shown in Fig. 3, semen parameters were superior in P-BS samples, reflecting changes as the reproductive season waned. P-BS diluted semen samples showed significantly higher progressive sperm motility (88.1±1.3% vs. 55.0±7.4%; P=0.0020), and trends toward higher sperm concentration (406±148 vs. 124±50 × 10⁶/mL; P= 0.1019) and normal sperm morphology (76.9±4.6% vs. 59.2±9.6%; P=0.0884). Sperm morphological defects that were more common in A-BS included significantly increased principal piece defects (1.2±0.4% vs. 7.8±4.8%; P=0.0140) and a trend toward elevated midpiece defects (10.9±4.1% vs. 25.2±6.3%; P=0.0764). No significant differences were found in the proportions of detached heads (4.4±0.7% vs. 6.2±1.7%; P=0.4296), cytoplasmic droplets (4.3±1.9% vs. 2.8±0.6%; P=0.7712), head defects (2.3±0.5% vs. 1.6±0.2%; P=0.4296), or acrosome defects (1.2±0.5% vs. 0.4±0.2%; P=0.4196). Additionally, P-BS diluted semen samples tended to have a higher total sperm count (203.2±74.2 vs. 62.2±24.9 × 10⁶; P=0.1019), with significantly more total progressively motile sperm (179.4±76.0 vs. 31.4±11.9 × 10⁶; P=0.0062) and total PMMN sperm (147.7±68.5 vs. 21.7±10.3 × 10⁶; P=0.0109) per ejaculate.

**Fig. 3.**
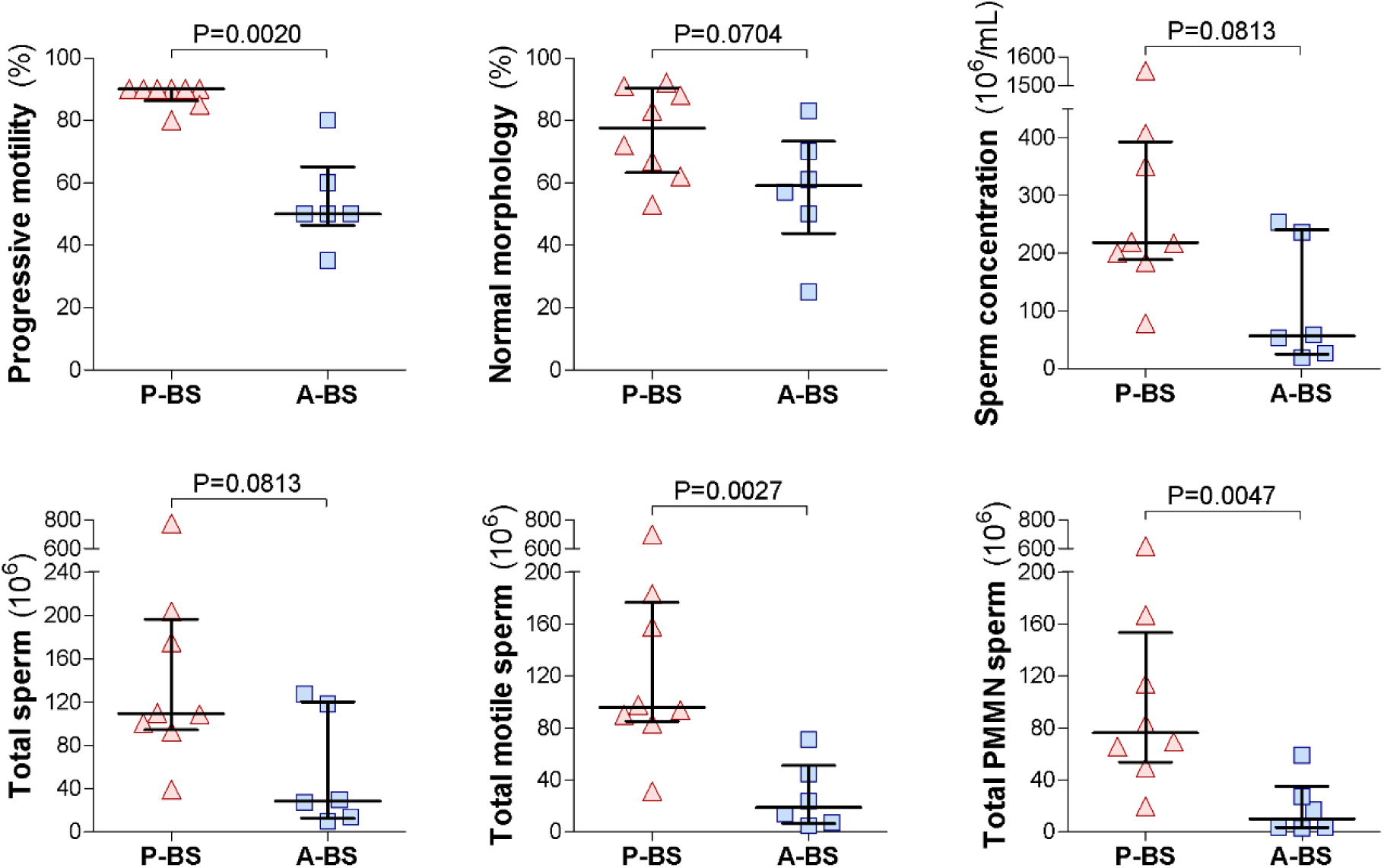
Seasonal variation in semen parameters of wild rock hyraxes collected via electroejaculation in 2021. Semen samples were obtained from wild males just before the onset of peak reproductive activity (P-BS) and again 3–4 weeks later (A-BS). Ejaculates were collected via electroejaculation directly into 500 µL of semen extender (EquiPlus, Minitube) for sperm preservation during the initial analysis. Based on the analyses of the extender-diluted semen samples, shown are progressive motility, normal morphology, sperm concentration, as well as total sperm per ejaculate, total motile sperm per ejaculate, and total progressively motile morphologically normal (PMMN) sperm per ejaculate. Each dot represents an individual extender-diluted semen sample; horizontal lines indicate the median, and whiskers show the interquartile range (IQR). Comparisons between P-BS and A-BS were made using the Mann–Whitney test, with P-values indicated in each panel.

### Electroejaculation-based semen collection and parameters in wild rock hyraxes – Summer of 2022

In the summer of 2022, 17 electroejaculation procedures were conducted: 11 during P-BS (July 1–7; n=9 males) and 6 during A-BS (August 4–9; n=6 males). Erection was achieved in all cases (100%), and semen was successfully collected in 15/17 attempts (88.2%). In P-BS, sperm cells were detected in 9/11 samples (81.8%), each from a different male. The two unsuccessful attempts included one pre-sperm sample and one contaminated by urine, a phenomenon reported in other species and attributed to bladder contraction induced by stimulation (Gomes-Alves, et al. 2014, O’Brien, et al. 2015, Zambelli and Cunto 2006); both males were recaptured and successfully sampled later in the session. In A-BS, semen was collected in all 6 procedures (100%) from six individual males.

Semen parameters from all 17 diluted semen samples collected during the summer of 2022 exhibited considerable variability in sperm quantity and quality. The numerical difference in progressive sperm motility between samples in P-BS versus A-BS was not statistically significant (P-BS: 80.6±6.9% vs. A-BS: 56.7±16.6%; P=0.6309). However, as illustrated in Fig. 4, all other semen parameters were notably superior in P-BS samples. The proportion of sperm with normal morphology was significantly higher in P-BS than in A-BS (82.4±2.7% vs. 66.2±4.0%; P=0.0048). Morphological defects more prevalent in A-BS included midpiece defects (7.0±1.1% vs. 13.8±1.1%; P=0.0018), distal cytoplasmic droplets (1.7±0.5% vs. 6.0±2.3%; P=0.0138), principal piece defects (0.9±0.3% vs. 3.2±0.6%; P=0.0030), acrosome defects (0.2±0.1% vs. 3.2±0.2%; P=0.0012), and head defects (0.7±0.2% vs. 2.8±1.0%; P=0.1183), but there was no difference in the incidence of detached heads between P-BS and A-BS (8.0±2.9% vs. 6.7 ±1.3%; P=0.7510). Sperm concentration tended to be higher in P-BS diluted semen samples (369±46 vs. 245±36 × 10⁶ sperm/mL; P=0.0879), as did the total sperm count per ejaculate (184±23 vs. 122.6±17.8 × 10⁶; P=0.0879). Notably, the total number of progressively motile sperm was significantly greater in P-BS (152.4±25.6 vs. 74.0±25.9 × 10⁶; P=0.0496), as was the total number of PMMN sperm per ejaculate (125.3±22.0 vs. 51.6±19.3 × 10⁶; P=0.0360).

**Fig. 4.**
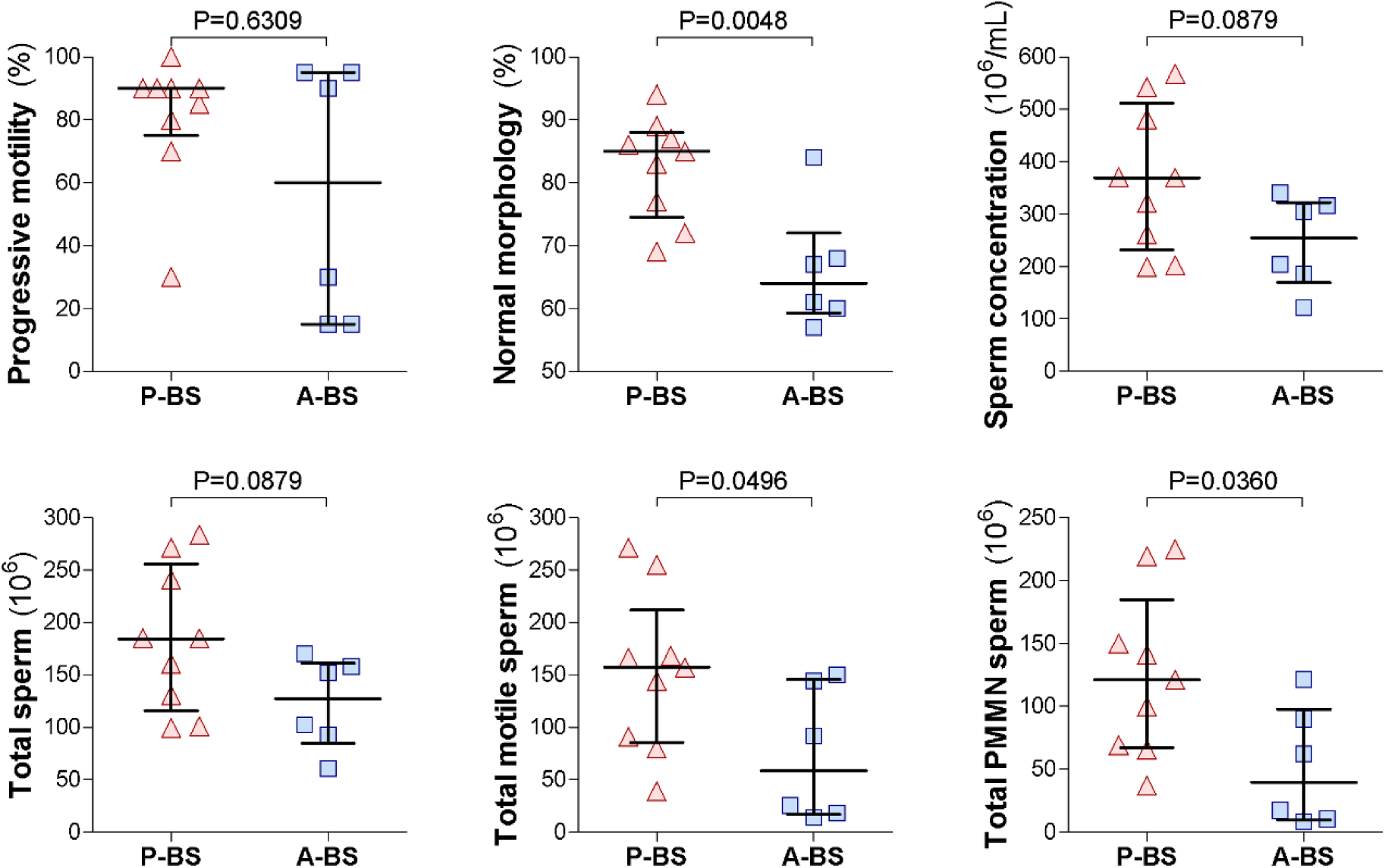
Seasonal variation in semen parameters of wild rock hyraxes collected via electroejaculation in 2022. Semen samples were obtained from wild males just before the onset of peak reproductive activity (P-BS) and again 3–4 weeks later (A-BS). Ejaculates were collected via electroejaculation directly into 500 µL of semen extender (EquiPlus, Minitube) for sperm preservation during the initial analysis. Based on the analyses of the extender-diluted semen samples, shown are progressive motility, normal morphology, sperm concentration, as well as total sperm per ejaculate, total motile sperm per ejaculate, and total progressively motile morphologically normal (PMMN) sperm per ejaculate. Each dot represents an individual extender-diluted semen sample; horizontal lines indicate the median, and whiskers show the interquartile range (IQR). Comparisons between P-BS and A-BS were made using the Mann–Whitney test, with P-values indicated in each panel.

### Electroejaculation-based semen collection and parameters in wild rock hyraxes – Summer of 2023

In the summer of 2023, 19 electroejaculation procedures were performed under field conditions: 9 during P-BS (n=8 males) and 10 during A-BS (n=10 males). Erection was achieved in all cases (100%), and sperm cells were collected in 18 of 19 ejaculates (94.7%) within a mean stimulation of 11±0.9 minutes (Median 10 min; range 5–20 min). The single unsuccessful attempt occurred during P-BS, but the male was recaptured the next day and successfully sampled. Semen parameters were recorded for all samples, except motility, which was missed in two cases (one per session) due to operator error.

Semen parameters from all 18 diluted semen samples collected during the summer of 2023 exhibited considerable variability in sperm quantity and quality. The percentage of sperm cells with progressive motility did not differ significantly between samples collected during the peak breeding season (P-BS; 87.1±11.2%) and those collected post-peak (A-BS; 84.7±6.1%; P=0.1564). However, consistent with findings from 2021 and 2022, semen parameters were generally superior in P-BS samples (Fig. 5). Sperm concentration was significantly higher in P-BS diluted semen samples than in A-BS (348±96 vs. 72±27 × 10⁶ sperm cells/mL; P=0.0164). Additionally, P-BS samples tended to have higher proportions of morphologically normal sperm (77.4±6% vs. 66.2±4%; P=0.0619). Morphological defect rates were as follows: midpiece defects (13.6±5.5% vs. 19.3±3.3%; P=0.0681), distal cytoplasmic droplets (3.7±1% vs. 3.0±1.3%; P=0.8926), principal piece defects (1.8±0.6% vs. 1.5±0.4%; P=0.8109), acrosome defects (0.9±0.6% vs. 0.8±0.2%; P=0.4669), head defects (1.9±0.8% vs. 2.4±0.4%; P=0.2321), and detached heads (4.1±0.7% vs. 5.4±0.8%; P=0.4393). On a per-ejaculate basis, P-BS diluted semen samples had significantly higher total sperm counts (all samples: 174±48 vs. 36±13.6 × 10⁶; P=0.0164; excluding two samples with missing motility data: 195±49 vs. 38.8±14.8 × 10⁶; P=0.0149), total progressively motile sperm cells (193±50 vs. 35.7±15.3 × 10⁶; P=0.0164), and total PMMN sperm (164±46 vs. 28.5±13.1 × 10⁶; P=0.0229).

**Fig. 5.**
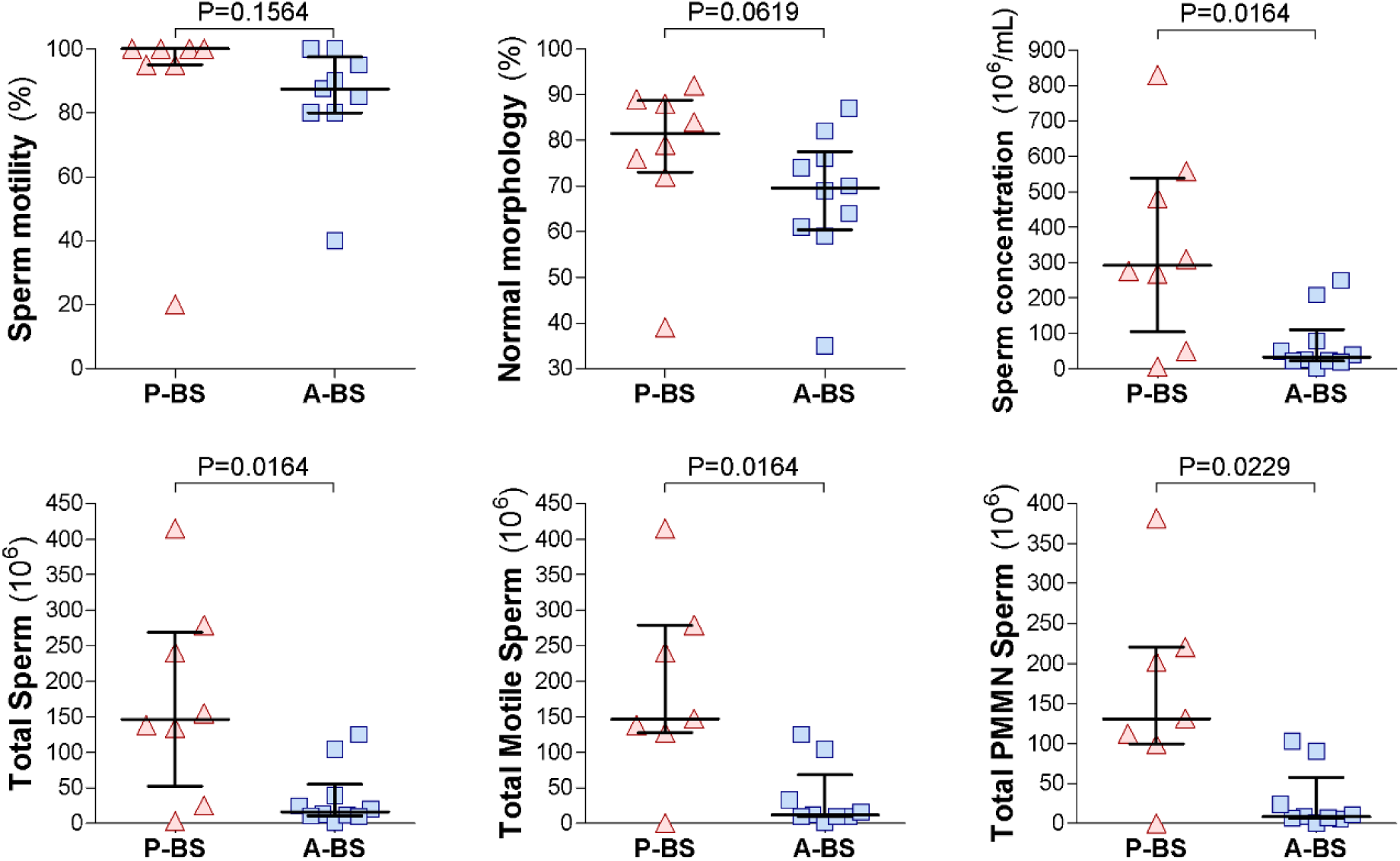
Seasonal variation in semen parameters of wild rock hyraxes collected via electroejaculation in 2023. Semen samples were obtained from wild males just before the onset of peak reproductive activity (P-BS) and again 3–4 weeks later (A-BS). Ejaculates were collected via electroejaculation directly into 500 µL of semen extender (EquiPlus, Minitube) for sperm preservation during initial analysis. Based on the analyses of the extender-diluted semen samples, shown are progressive motility, normal morphology, sperm concentration, total sperm per ejaculate, total motile sperm per ejaculate, and total progressively motile morphologically normal (PMMN) sperm per ejaculate. Each dot represents an individual extender-diluted semen sample; horizontal lines indicate the median, and whiskers show the inter-quartile range (IQR). Comparisons between P-BS and A-BS were made using the Mann–Whitney test, with P-values shown in each panel.

### Seasonal variation in semen parameters: repeated measures analysis across three breeding seasons (2021–2023)

Electroejaculation consistently induced synchronized muscle contractions and erections in all wild males sampled during the 2021–2023 breeding seasons. Sperm-containing samples were successfully obtained in 47 of 53 procedures (88.7%). All six failed attempts were followed by successful collections from the same males later in the sampling session. Our analysis revealed a higher success rate in A-BS (21/21) compared to P-BS (26/32), indicating a non-significant trend (P=0.0702). These findings confirm the overall reliability of electroejaculation in wild rock hyraxes.

To avoid potential bias stemming from differences in individual capture probability or inherent variability in semen characteristics, repeated measures ANOVA analyses were conducted using data from males sampled twice per breeding season in the same year, once in P-BS and once in A-BS (2021: n=4; 2022: n=6; 2023: n=5; total ejaculates=32). No significant differences were found in electroejaculation duration between sampling sessions (P-BS vs. A-BS, P=0.2152), years (P=0.6898), or their interaction (P=0.4571). However, all semen parameters were superior in P-BS diluted semen samples, as shown in Fig. 6 and Table 3, with no significant year effects or session × year interaction. The percentage of sperm cells with progressive motility was significantly higher during P-BS than A-BS (86.1±4.9% vs. 63.4±8.2%; P=0.0082), reflecting a mean reduction of 23.1±7.2% as the breeding season faded. Similarly, the mean sperm concentration was significantly higher in P-BS diluted semen samples (337.0±53.5 × 10⁶/mL vs. 163.0±30.5 × 10⁶/mL; P=0.0081), representing a reduction of 199.2±61.8 × 10⁶/mL. The percentage of sperm cells with normal morphology also declined significantly from P-BS to A-BS (79.3±3.9% vs. 64.8±3.7%; P=0.0076), with a mean reduction of 17.2±5.3%. Differential morphology analysis revealed no significant differences in the proportions of acrosome defects, head defects, detached heads, or cytoplasmic droplets. However, tendencies were observed for increased midpiece defects (P=0.065) and principal piece defects (P=0.0641) in A-BS samples. Round cells were rarely observed in either P-BS or A-BS samples (typically 0–3 per 100 spermatozoa), and thus were not further analyzed.

**Fig. 6.**
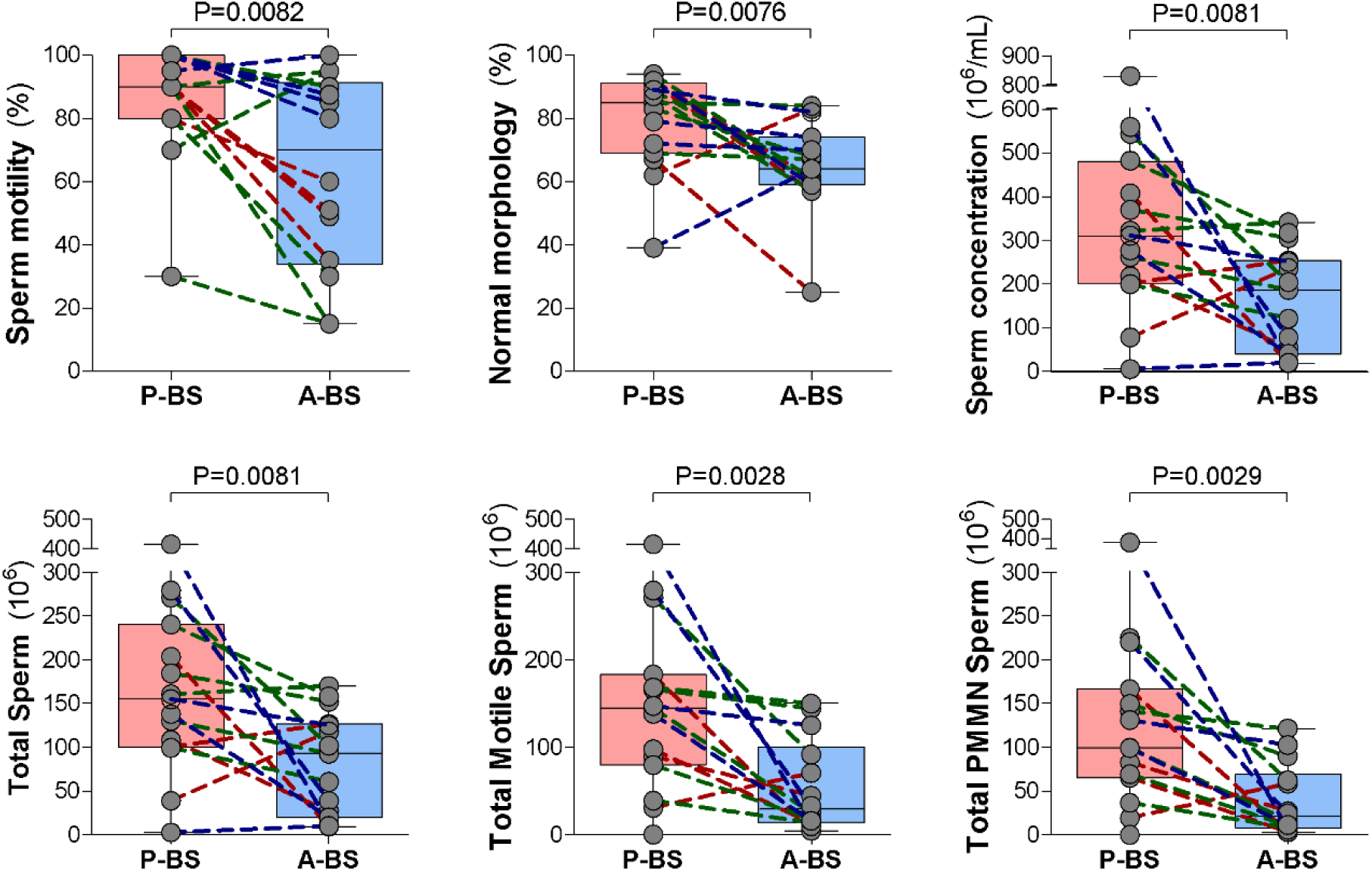
Repeated measures analysis of semen parameters in rock hyraxes sampled during and after the breeding season. Semen samples were collected from the same males twice within the same breeding season: once during peak breeding season (P-BS, late June–early July) and once approximately three weeks later (A-BS, late July–early August), in 2021 (n=4 males), 2022 (n=6 males), and 2023 (n=5 males). Repeated measures ANOVA analyses were conducted to evaluate the effects of sampling session (P-BS vs. A-BS), year of sampling (2021, 2022, 2023), and their interaction on the parameters assessed for extender-diluted semen samples. (A) Progressive motility (%) decreased significantly from P-BS to A-BS (Sampling session: P=0.0082; Year: P=0.1151; Sampling session × Year: P=0.3411). (B) Sperm concentration (×10^6 sperm cells/mL) was lower in A-BS (Sampling session: P=0.0081; Year: P=0.1233; Sampling session × Year: P=0.1380). (C) The proportion of morphologically normal sperm (%) declined significantly post-breeding season (Sampling session: P=0.0076; Year: P=0.2786; Sampling session × Year: P=0.7709). (D) Total sperm count per extender-diluted semen sample (×10^6) was significantly reduced in A-BS (Sampling session: P=0.0081; Year: P=0.1233; Sampling session × Year: P=0.1380). (E) The number of progressively motile sperm cells per extender-diluted semen sample (×10^6) also declined (Sampling session: P=0.0028; Year: P=0.1611; Sampling session × Year: P=0.1735). (F) The total number of progressively motile, morphologically normal (PMMN) sperm cells per ejaculate (×10^6) was significantly lower post-breeding season (Sampling session: P=0.0029; Year: P=0.1667; Sampling session × Year: P=0.2214). Each panel shows individual semen samples overlaid on box-and-whisker plots (line in the middle=median; box=interquartile range; whiskers=minimum and maximum). Paired samples from the same male are connected by dotted lines, colored by year (red for 2021, green for 2022, blue for 2023).

**Table 3.** Seasonal changes in semen parameters in rock hyraxes sampled during the peak breeding season (P-BS) and again approximately 3–4 weeks later (A-BS) across three breeding seasons (2021–2023)

| Parameter | P-BS<sup>1</sup> | A-BS<sup>1</sup> | P-value<sup>2</sup> |
| --- | --- | --- | --- |
| Body weight (kg) | 2.77±0.13<br>2.75 (2.3–3.25) | 2.62±0.11<br>2.60 (2.3–3) | 0.0022 |
| Electroejaculation duration (min) | 11.9±1.6<br>11.0 (8–16) | 10.0±0.9<br>10.0 (8–12) | 0.2152 |
| Progressive sperm motility (%) | 86.1±4.9<br>90 (80–100) | 63.4±8.2<br>70 (33.8–91.3) | 0.0082 |
| Sperm concentration (x 10<sup>6</sup>/mL) | 337.0±53.5<br>310 (200–481) | 163.00±30.5<br>186 (40–254) | 0.0081 |
| **Sperm morphology** |  |  |  |
| Normal morphology (%) | 79.3±3.9<br>85 (69–91) | 64.8±3.7<br>64 (59–74) | 0.0076 |
| Acrosome defects (%) | 0.6±0.3<br>0 (0–1) | 1.7±0.5<br>1 (0–3) | 0.1418 |
| Head defects (%) | 1.3±0.4<br>1 (0–2) | 2.3±0.5<br>2 (1–4) | 0.1739 |
| Midpiece defects (%) | 11.1±3.3<br>8 (3–14) | 18.7±2.5<br>16 (12–22) | 0.065 |
| Principal piece defects (%) | 1.3±0.4<br>1 (1–1) | 4.2±1.7<br>3 (2–4) | 0.0641 |
| Detached heads (%) | 5.4±1.8<br>5 (2–6) | 6.1±0.9<br>6 (3–9) | 0.6875 |
| Cytoplasmic droplets (%) | 3.2±0.8<br>3 (1–4) | 4.0±1.0<br>3 (2–4) | 0.7515 |
| Total sperm (x 10<sup>6</sup>) | 168.5±26.7<br>155 (100–240.5) | 81.5±15.2<br>93 (20–127) | 0.0081 |
| Total motile sperm (x 10<sup>6</sup>) | 160.7±27.7<br>145.9 (87.4–205.1) | 54.4±14.0<br>29.3 (13.8–100.1) | 0.0028 |
| Total PMMN sperm (X10<sup>6</sup>) | 134.9±25.1<br>115.4 (68.3–180.1) | 38.9±10.8<br>20.8 (7.9–69.3) | 0.0022 |

<sup>1</sup> Values are presented as mean±SEM (top row) and median (interquartile range) (bottom row).
<sup>2</sup> The P-values were calculated using repeated measures ANOVA, including only males sampled at both time points within the same year (2021, 2022, or 2023). No significant effects of year or year × sampling session interactions were observed and are therefore not shown.

Total sperm output and functional sperm subpopulations were also significantly lower during A-BS. The total sperm count per diluted semen sample decreased significantly from P-BS to A-BS (168.5±26.7 × 10⁶ vs. 81.5±15.2 × 10⁶; P=0.0081), with a mean reduction of 99.6±30.9 × 10⁶ sperm cells per ejaculate. The total number of motile sperm declined by 112.2±29.3 × 10⁶ sperm cells (160.7±27.7 × 10⁶ vs. 54.4±14.0 × 10⁶; P=0.0028), and the total number of PMMN sperm decreased by 100.5±26.4 × 10⁶ (134.9±25.1 × 10⁶ vs. 38.9±10.8 × 10⁶; P=0.0022). Interestingly, body weight was significantly higher in males during P-BS compared to A-BS (2.77±0.13 kg vs. 2.62±0.11 kg; P=0.0022), with a mean loss of 145±38 grams, potentially reflecting a seasonal shift in physiological condition.

## Discussion

In this study, we successfully developed and implemented a non-terminal electroejaculation protocol for semen collection in the rock hyrax, enabling for the first time comprehensive in vivo assessments of sperm morphology, ultrastructure, and morphometry, as well as detailed evaluations of semen quality and its seasonal variation in both captive and wild populations. The procedure was shown to be safe, practical, and well-tolerated in laboratory and field settings, eliciting consistent physiological responses such as synchronized muscle contractions and reliable erections in all males. Sperm-containing ejaculates were obtained at high success rates, particularly around the window of seasonal reproductive activity. Ejaculation was achieved within 8–12 minutes of stimulation in most males (range: 5–20 minutes), using voltage levels comparable to or lower than those used in humans and other wild and domestic species (Abaigar, et al. 2001, Chung, et al. 1995, Okano, et al. 2006, Takaesu, et al. 2022). Although ejaculate volumes were small, the milky opaque appearance consistently indicated sperm presence, compared to clear or yellowish fluids, which did not contain sperm. Importantly, no adverse effects were observed, reinforcing electroejaculation as a safe and repeatable tool for longitudinal reproductive and conservation-oriented studies in rock hyraxes.

The structural organization of rock hyrax spermatozoa comprises a compact head, midpiece, and an elongated principal piece (Bedford and Millar 1978). High-resolution SEM imaging confirmed this architecture, as well as revealed intact acrosomal, post-acrosomal membranes and a defined equatorial segment. Furthermore, the detected ultrastructural organization of hyrax sperm, including a well-defined axoneme, an elongated principal piece, and a mitochondria-rich midpiece with helically arranged mitochondria, reflects motility-enhancing adaptations commonly observed among eutherian mammals, including species from Afrotheria and Laurasiatheria, supporting efficient progressive motility propulsion through viscous environments such as the female reproductive tract (Bedford and Millar 1978, Gu, et al. 2019, Korneev, et al. 2021, Thompson, et al. 2018). Our morphometric analysis revealed that in *Procavia capensis*, the average sperm length is ∼56 µm, with the flagellum accounting for ∼91% of this length (∼18% midpiece, ∼73% principal piece). This proportion aligns with or slightly exceeds values reported by Gu et al. (2019) for other non-rodent mammalian species, including humans (∼92%), dogs (∼91%), bulls (∼88%), goat (87%), rabbits (86%), and pigs (∼86%). The relatively high flagellum-to-head ratio and head compactness in hyrax sperm may reflect species-specific adaptations to enhance sperm velocity, possibly linked to selective pressures such as sperm competition and constrained mating seasons (Fisher, et al. 2013, Gomendio and Roldan 1991, Gu, et al. 2019, Thompson, et al. 2018). Although the rock hyrax exhibits a predominantly polygynous mating system, opportunities for postcopulatory sperm competition may occur due to overlapping breeding activity among males. Therefore, sperm competition should be considered as a potential, albeit unconfirmed, selective pressure contributing to the evolution of these sperm traits.

In addition to normal sperm forms, morphological evaluations based on N&E-stained slides and SEM revealed a broad spectrum of abnormalities, which were more common in samples collected after the peak breeding season. Defects affecting the midpiece and principal piece were most frequent and included anomalies such as distal midpiece reflexes, mitochondrial sheath disorganization, coiled or truncated tails, and double axonemal granules. These regions are critical for sperm motility and energy production, and damage to their structure likely reflects disrupted spermiogenesis or oxidative stress during the seasonal transition out of reproductive activity (Pintus, et al. 2015, Schmoll, et al. 2018). SEM images revealed multiple structural abnormalities, including cytoplasmic droplets in both proximal and distal locations, torn fibrous sheaths at the annulus, exposed axonemal microtubules, and detached heads often accompanied by nuclear membrane rupture and possible chromatin leakage. These features may reflect incomplete epididymal maturation, compromised sperm integrity, or post-ejaculatory degradation.

Cytological analyses provided additional insight into testicular output and germ cell quality. N&E staining enabled efficient morphological screening, while Diff-Quick staining improved nuclear and cytoplasmic resolution, allowing the identification of specific round cells, including presumptive spermatocytes, round spermatids, and degenerative medusa cells. In captive males, round cells were particularly abundant in samples collected farther from the breeding season, and in association with low-quality semen, supporting seasonal testicular regression and impaired spermatogenesis (Beltran-Frutos, et al. 2022, Johanisson, et al. 2000). Their higher prevalence in captive males likely reflects broader year-round sampling, in contrast to wild males sampled only near peak fertility; however, differences between captive and wild habitats may have also contributed to this disparity. Although detached heads and cytoplasmic droplets did not differ significantly between time points, their persistence, especially post-peak, suggests ongoing disruption in epididymal processing. Together, SEM and cytological analyses provided a detailed and complementary view of sperm development and integrity, revealing seasonal degeneration patterns that align with the rock hyrax’s sharply timed breeding behavior.

Seasonal changes in semen quality were consistent across all three years of field sampling. Semen collected during the peak breeding season showed significantly higher sperm concentration, progressive motility, and normal morphology compared to samples collected 3–4 weeks later. Consistent with the notion that reproductive seasonality and sperm competition influence ejaculate traits (Thompson, et al. 2018), our findings on the marked post-peak declines in sperm quality and structure, as overall illustrated in the steep reduction in motile and PMMN sperm subpopulations, suggest constrained temporal windows for optimal male fertility. Moreover, the subsequent decline parallels that seen in other seasonal breeders, reflecting reduced testicular function and hormone fluctuations that impair spermatogenesis and germ cell integrity, factors that likely contributed to the elevated round cell counts observed post-peak in our hyrax study (Beltran-Frutos, et al. 2022, Goeritz, et al. 2003, Huamani, et al. 2025). Notably, in addition to changes in semen quality, a significant seasonal decline in body weight was observed in wild hyrax males P-BS to A-BS. This may reflect a shift in energy allocation away from reproduction as the season ends, a pattern also seen in other seasonal breeders such as the eastern spotted skunk (*Spilogale putorius ambarvilus*) (Kaplan and Mead 1994), masked palm civet (*Paguma larvata*) (Tsui, et al. 1974), and Vespertilionid Bat (*Myotis nigricans*) (Beguelini, et al. 2015), where reductions in androgen production and testicular activity are accompanied by decreases in body condition. Alternatively, weight loss may result from the high energetic demands of reproductive activity, such as increased movement, vocalizations, and competition, which intensify during the mating season in rock hyraxes (Goll, et al. 2023).

As in many wildlife reproduction studies, several methodological and biological limitations should be acknowledged. Although electroejaculation proved safe and effective in the rock hyrax, animal welfare concerns have been raised regarding its use in other species, mainly due to potential discomfort during stimulation (Abril-Sanchez, et al. 2019, Palmer 2005). In our study, all procedures were performed under anesthesia, and no adverse effects were observed, minimizing welfare risks. Trapping and handling times were kept as short as possible to reduce stress, and animals were monitored until they fully recovered before being released at their original capture sites. Consistent with this cautious approach, in captive males, stimulation duration was initially limited to 10 minutes as a precaution, since this was the first application of electroejaculation in this species; after confirming its safety, the duration was extended to up to 20 minutes for wild collections to increase success rates. It should also be noted that electroejaculated semen may not fully reflect the composition of naturally ejaculated semen, as differences in spermatozoa and seminal plasma content and glandular stimulation can occur (Lv, et al. 2025, Marco-Jimenez, et al. 2005, Zambelli and Cunto 2006). In addition, semen collection conditions, including the choice of extender, dilutions, handling time, and temperature control, may have influenced some of the measured parameters, and optimization of these factors should be considered in future studies. Moreover, our semen evaluation relied on manual methods, which can introduce a degree of subjectivity; however, all analyses, except for motility assessment, were performed blindly by a single trained operator to ensure consistency. Computer-Aided Sperm Analysis (CASA) could provide more detailed and objective assessments of sperm motility and kinematics (Brito 2025, van der Horst 2020, 2021); however, this technique was not feasible under our remote field conditions, where all equipment had to be transported on foot and powered by batteries, and was unavailable at Bar-Ilan University during captive sampling. Future studies could benefit from incorporating CASA when appropriate laboratory facilities are available. Finally, potential effects of social dominance and sperm competition were not evaluated in our current analyses and represent an important avenue for future research (Bartlett, et al. 2017, Tamara Montrose, et al. 2008).

Overall, our findings demonstrate that male rock hyraxes exhibit marked seasonality in reproductive function, with semen quality, sperm structure, body condition, and reproductive behavior tightly aligned with the narrow mating window. While previous studies have primarily focused on behavioral aspects of reproduction, our work adds critical physiological insight by characterizing semen quality and its seasonal dynamics. The consistent decline in sperm quality after the peak breeding period, observed across years and individuals, underscores the importance of precise timing in fertility assessments and reproductive studies. The electroejaculation protocol established here offers a valuable, repeatable, and non-lethal tool for monitoring male reproductive status in rock hyraxes and may also be adapted for use in future longitudinal studies, in rock hyraxes and perhaps in other similarly sized wild mammals with seasonal breeding patterns or conservation relevance.

## Declarations

### Contributions

T.R. conceived and supervised the study, instructed semen collection, analyzed all laboratory samples and data, and wrote the manuscript. H.W. captured and performed semen collection in wild males and handled initial field processing. S.A.S. conducted semen collection from captive hyraxes and initial analysis. N.S. assisted with captive collections and performed morphometric measurements. N.W. provided technical support throughout and assisted with SEM analysis. L.K. co-conceived and supervised the study. All authors read, revised, and eventually approved the final manuscript.

## Acknowledgments

The authors thank Dr. Einat Zelinger of the Interdepartmental Equipment Facility, Faculty of Agriculture, Food and Environmental Sciences, The Hebrew University, for her assistance with the SEM analysis. We are also grateful to Dr. Shaked Druker from the Raz laboratory for his help with the electroejaculation equipment.

## References

Abaigar, T, M Cano, AR Pickard, and WV Holt 2001 Use of computer-assisted sperm motility assessment and multivariate pattern analysis to characterize ejaculate quality in Mohor gazelles (Gazella dama mhorr): effects of body weight, electroejaculation technique and short-term semen storage. Reproduction 122 265–273.

Abril-Sanchez, S, A Freitas-de-Melo, J Giriboni, J Santiago-Moreno, and R Ungerfeld 2019 Sperm collection by electroejaculation in small ruminants: A review on welfare problems and alternative techniques. Anim Reprod Sci 205 1–9.

Bar Ziv, E, A Ilany, V Demartsev, A Barocas, E Geffen, and L Koren 2016 Individual, social, and sexual niche traits affect copulation success in a polygynandrous mating system. Behavioral Ecology and Sociobiology 70 901–912.

Barocas, A, A Ilany, L Koren, M Kam, and E Geffen 2011 Variance in centrality within rock hyrax social networks predicts adult longevity. PLoS One 6 e22375.

Barth, AD, and RJ Oko 1989 Abnormal morphology of bovine spermatozoa / A.D. Barth, R.J. Oko. edn 1st ed. Ames: Iowa State University Press.

Bartlett, MJ, TE Steeves, NJ Gemmell, and PC Rosengrave 2017 Sperm competition risk drives rapid ejaculate adjustments mediated by seminal fluid. Elife 6.

Bedford, JM, and RP Millar 1978 The character of sperm maturation in the epididymis of the Ascrotal hyrax, Procavia capensis and armadillo, Dasypus novemcinctus. Biol Reprod 19 396–406.

Beguelini, MR, RM Goes, P Rahal, E Morielle-Versute, and SR Taboga 2015 Impact of the processes of testicular regression and recrudescence in the prostatic complex of the bat Myotis nigricans (Chiroptera: Vespertilionidae). J Morphol 276 721–732.

Beltran-Frutos, E, V Seco-Rovira, J Martinez-Hernandez, C Ferrer, MI Serrano-Sanchez, and LM Pastor 2022 Cellular Modifications in Spermatogenesis during Seasonal Testicular Regression: An Update Review in Mammals. Animals (Basel) 12.

Bordes, CNM, R Beukeboom, Y Goll, L Koren, and A Ilany 2022 High-resolution tracking of hyrax social interactions highlights nighttime drivers of animal sociality. Commun Biol 5 1378.

Brito, LF, LM Greene, A Kelleman, M Knobbe, and R Turner 2011 Effect of method and clinician on stallion sperm morphology evaluation. Theriogenology 76 745–750.

Brito, LF, A Kelleman, LM Greene, T Raz, and AD Barth 2010 Semen characteristics in a sub-fertile Arabian stallion with idiopathic teratospermia. Reprod Domest Anim 45 354–358.

Brito, LFC 2025 Sperm Motility Evaluation in Stallion Fresh, Cooled and Frozen Semen Using a Portable Computer-Assisted Sperm Analysis System. Reprod Domest Anim 60 e70052.

Brito, LFC, GC Althouse, C Aurich, PJ Chenoweth, BE Eilts, CC Love, GC Luvoni, JR Mitchell, AT Peter, DG Pugh, and D Waberski 2016 Andrology laboratory review: Evaluation of sperm concentration. Theriogenology 85 1507–1527.

Cary, JA, S Madill, K Farnsworth, JT Hayna, L Duoos, and ML Fahning 2004 A comparison of electroejaculation and epididymal sperm collection techniques in stallions. Can Vet J 45 35–41.

Chung, PH, G Palermo, PN Schlegel, LL Veeck, JF Eid, and Z Rosenwaks 1998 The use of intracytoplasmic sperm injection with electroejaculates from anejaculatory men. Hum Reprod 13 1854–1858.

Chung, PH, TR Yeko, JC Mayer, EJ Sanford, and GB Maroulis 1995 Assisted fertility using electroejaculation in men with spinal cord injury--a review of literature. Fertil Steril 64 1–9.

Demartsev, V, M Haddas-Sasson, A Ilany, L Koren, and E Geffen 2023 Male rock hyraxes that maintain an isochronous song rhythm achieve higher reproductive success. J Anim Ecol 92 1520–1531.

Fisher, DO, CR Dickman, ME Jones, and SP Blomberg 2013 Sperm competition drives the evolution of suicidal reproduction in mammals. Proc Natl Acad Sci U S A 110 17910–17914.

Glover, TD 1973 Aspects of sperm production in some East African mammals. J Reprod Fertil 35 45–53.

Glover, TD, and JB Sale 1968 Reproductive system of male rock hyrax (procavia and heterohyrax) Journal of Zoology 156 351-&.

Goeritz, F, M Quest, A Wagener, M Fassbender, A Broich, TB Hildebrandt, RR Hofmann, and S Blottner 2003 Seasonal timing of sperm production in roe deer: interrelationship among changes in ejaculate parameters, morphology and function of testis and accessory glands. Theriogenology 59 1487–1502.

Goll, Y, C Bordes, YA Weissman, I Shnitzer, R Beukeboom, A Ilany, L Koren, and E Geffen 2023 The interaction between cortisol and testosterone predicts leadership within rock hyrax social groups. Sci Rep 13 14857.

Gomendio, M, and ER Roldan 1991 Sperm competition influences sperm size in mammals. Proc Biol Sci 243 181–185.

Gomes-Alves, S, M Alvarez, M Nicolas, C Martinez-Rodriguez, S Borragan, CA Chamorro, L Anel, and P de Paz 2014 Salvaging urospermic ejaculates from brown bear (Ursus arctos). Anim Reprod Sci 150 148–157.

Gu, NH, WL Zhao, GS Wang, and F Sun 2019 Comparative analysis of mammalian sperm ultrastructure reveals relationships between sperm morphology, mitochondrial functions and motility. Reprod Biol Endocrinol 17 66.

Huamani, MC, CYG Palomino, HRO Berrocal, IML Arcce, DK Bellido-Quispe, AYA Leiva, MS Chaves, and VJ de Figueiredo Freitas 2025 Influence of reproductive season and testicular volume on seminal parameters of alpacas (Vicugna pacos). Trop Anim Health Prod 57 227.

Johanisson, E, A Campana, R Luthi, and A de Agostini 2000 Evaluation of ‘round cells’ in semen analysis: a comparative study. Hum Reprod Update 6 404–412.

Kaplan, JB, and RA Mead 1994 Seasonal Changes in Testicular Function and Seminal Characteristics of the Male Eastern Spotted Skunk (Spilogale putorius ambarvilus). Journal of Mammalogy 75 1013–1020.

Koren, L, O Mokady, and E Geffen 2006 Elevated testosterone levels and social ranks in female rock hyrax. Horm Behav 49 470–477.

Koren, L, O Mokady, and E Geffen 2008 Social status and cortisol levels in singing rock hyraxes. Horm Behav 54 212–216.

Koresh, E, D Matas, and L Koren 2016 Experimental elevation of wildlife testosterone using silastic tube implants. Res Vet Sci 108 1–7.

Korneev, D, DJ Merriner, G Gervinskas, A de Marco, and MK O’Bryan 2021 New Insights Into Sperm Ultrastructure Through Enhanced Scanning Electron Microscopy. Front Cell Dev Biol 9 672592.

Koziol, JH, and CL Armstrong 2022 Normal Detached Heads and Free Abnormal Heads, Sperm Morphology of Domestic Animals, pp. 44–46.

Lv, C, A Larbi, J Liang, C Li, B Bouabid, G Wu, and G Quan 2025 Effects of semen collection methods on sperm quality and metabolite profile in goat seminal plasma: Comparing between artificial vagina and electro-ejaculator techniques. Anim Reprod Sci 279 107885.

Marco-Jimenez, F, S Puchades, J Gadea, JS Vicente, and MP Viudes-de-Castro 2005 Effect of semen collection method on pre- and post-thaw Guirra ram spermatozoa. Theriogenology 64 1756–1765.

Millar, RP 1972 Degradation of spermatozoa in the epididymis of a seasonally breeding mammal, the rock hyrax, Procavia capensis. J Reprod Fertil 30 447–450.

Millar, RP, and TD Glover 1970 Seasonal changes in reproductive tract of male rock hyrax, Procavia capensis. Journal of Reproduction and Fertility 23 497-&.

O’Brien, JK, TL Roth, MA Stoops, RL Ball, KJ Steinman, GA Montano, CC Love, and TR Robeck 2015 Sperm sex-sorting and preservation for managing the sex ratio and genetic diversity of the southern white rhinoceros (Ceratotherium simum simum). Anim Reprod Sci 152 137–153.

Okano, T, T Murase, S Nakamura, T Komatsu, T Tsubota, and M Asano 2009 Normal sperm morphology and changes of semen characteristics and abnormal morphological spermatozoa among peri-mating seasons in captive japanese black bears (Ursus thibetanus japonicus). J Reprod Dev 55 194–199.

Okano, T, T Murase, C Yayota, T Komatsu, K Miyazawa, M Asano, and T Tsubota 2006 Characteristics of captive Japanese black bears (Ursus thibetanus japonicus) semen collected by electroejaculation with different voltages for stimulation and frozen-thawed under different conditions. Anim Reprod Sci 95 134–143.

Palmer, CW 2005 Welfare aspects of theriogenology: investigating alternatives to electroejaculation of bulls. Theriogenology 64 469–479.

Pintus, E, JL Ros-Santaella, and JJ Garde 2015 Beyond Testis Size: Links between Spermatogenesis and Sperm Traits in a Seasonal Breeding Mammal. PLoS One 10 e0139240.

Sale, JB 1965 Gestation Period and Neonatal Weight of the Hyrax. Nature 205 1240–1241.

Schmoll, T, O Kleven, and M Rusche 2018 Individual phenotypic plasticity explains seasonal variation in sperm morphology in a passerine bird. Evolutionary Ecology Research 19 547–560.

Setchell, BP 1993 Male reproduction. Reproduction in Domesticated Animals 83–128.

.Soma, H, T Spraker, and K Benirschke 1976 Macerated Monozygotic Twins in Quadruplet Pregnancy of the Cape hyrax (*Procavia capensis*). Journal of the Mammalogical Society of Japan 6 210–213.

Takaesu, N, C Kanno, K Sugimoto, M Nagano, A Kaneko, Y Indo, H Imai, H Hirai, M Okamoto, M Sashika, M Shimozuru, S Katagiri, T Tsubota, and Y Yanagawa 2022 Semen collection by urethral catheterization and electro-ejaculation with different voltages, and the effect of holding temperature and cooling rate before cryopreservation on semen quality in the Japanese macaque (Macaca fuscata). J Vet Med Sci 84 429–438.

Tamara Montrose, V, W Edwin Harris, AJ Moore, and PJ Moore 2008 Sperm competition within a dominance hierarchy: investment in social status vs. investment in ejaculates. J Evol Biol 21 1290–1296.

Tecirlioglu, RT, ES Hayes, and AO Trounson 2002 Semen collection from mice: electroejaculation. Reprod Fertil Dev 14 363–371.

Thompson, SK, NA Kutchy, S Kwok, ZNA Rosyada, IG Imumorin, B Purwantara, and E Memili 2018 Review: Sperm: Comparative morphology and function related to altered reproductive strategies and fertility in mammals. The Professional Animal Scientist 34 558–565.

Tsui, HW, WH Tam, B Lofts, and JG Phillips 1974 The annual testicular cycle and androgen production in vitro in the masked civet cat, Paguma l. larvata. J Reprod Fertil 36 283–293.

van der Horst, G 2020 Computer Aided Sperm Analysis (CASA) in domestic animals: Current status, three D tracking and flagellar analysis. Anim Reprod Sci 220 106350.

van der Horst, G 2021 Status of Sperm Functionality Assessment in Wildlife Species: From Fish to Primates. Animals (Basel) 11.

Weissman, YA, V Demartsev, A Ilany, A Barocas, E Bar-Ziv, E Geffen, and L Koren 2019 Social context mediates testosterone’s effect on snort acoustics in male hyrax songs. Horm Behav 114 104535.

World Health Organization 2021 WHO laboratory manual for the examination and processing of human semen. edn 6th. Geneva: World Health Organization.

Zambelli, D, and M Cunto 2006 Semen collection in cats: techniques and analysis. Theriogenology 66 159–165.

