## Supplementary material for "In vivo semen characterization and seasonal variation in *Procavia capensis*": Supp Table 1

**Table S1.** Summary of semen collection attempts and responses to electroejaculation in captive rock hyrax males (September 2017 - October 2018).

| No. | Date | Male ID | BW (kg) | Erection: Yes/No (stimulation parameters) | Ejaculation: Yes/No | Ejaculation: Physical appearance | Ejaculation: Presence of Sperm Cells (Yes/No) |
|---|---|---|---|---|---|---|---|
| 1 | 17 Sep 2017 | C3 | 3.3 | Yes | Yes | Milky opaque | Yes |
| 2 | 16 Nov 2017 | C1 | 2.5 | Yes | Yes | Clear | No (Only round cells) |
| 3 | 16 Nov 2017 | C2 | 2.6 | Yes | No | --- | --- |
| 4 | 28 Jan 2018 | C1 | 2.6 | Yes | No | --- | --- |
| 5 | 28 Jan 2018 | C2 | 2.6 | Yes | Yes | Clear yellowish | No |
| 6 | 28 Mar 2018 | C1 | 3.1 | Yes (within ~7 min) | No | --- | --- |
| 7 | 28 Mar 2018 | C2 | 2.7 | Yes (within ~3 min) | No | --- | --- |
| 8 | 29 Apr 2018 | C1 | 3.2 | Yes (within ~2.5 min) | Yes | Clear yellowish, then viscous | No |
| 9 | 29 Apr 2018 | C2 | 2.8 | Yes (within ~2.5 min; at 4V) | Yes | Clear yellowish, then viscous | No |
| 10 | 16 May 2018 | C1 | 3.3 | Yes (within ~2 min; at 4V) | Yes | Clear-milky | No (Only round cells) |
| 11 | 16 May 2018 | C2 | 2.8 | Yes (within ~2 min; at 3V) | Yes | Clear-milky | No (Only round cells) |
| 12 | 19 Jul 2018 | C2 | 2.9 | Yes (within ~3.5 min; at 3.5V) | Yes | Milky opaque | Yes |
| 13 | 12 Aug 2018 | C1 | 3.1 | Yes (within ~8.5 min; at 8.5V) | Yes | Milky opaque | Yes |
| 14 | 13 Sep 2018 | C2 | 2.9 | Yes (within ~6.5 min; at 7V) | Yes | Milky opaque | Yes |
| 15 | 17 Sep 2018 | C2 | 3 | Yes (within ~2 min; at 3V) | Yes | Milky opaque | Yes |
| 16 | 9 Oct 2018 | C2 | 3.2 | Yes (Partial) (within ~6.5 min; at 4V) | No | --- | --- |
| 17 | 9 Oct 2018 | C1 | 3.1 | Yes (within ~6 min; at 6V) | Yes | Milky opaque | Yes |
